# Identifying a supramodal language network in human brain with individual fingerprint

**DOI:** 10.1101/2020.05.10.085787

**Authors:** Lanfang Liu, Xin Yan, Hehui Li, Dingguo Gao, Guosheng Ding

**Affiliations:** Guangdong Provincial Key Laboratory of Social Cognitive Neuroscience and Mental Health, Department of Psychology, Sun Yat-sen University, Guangzhou 510006, China; State Key Laboratory of Cognitive Neuroscience and Learning, Beijing Normal University & IDG/McGovern Institute for Brain Research, Beijing, 100875, China; Department of Communicative Sciences and Disorders, Michigan State University, East Lansing Michigan, 48823, United States; Mental Health Center, Wenhua College, Wuhan, 430000, China

**Keywords:** language comprehension, supramodal, fingerprint, fMRI

## Abstract

Where is human language processed in the brain independent of its form? We addressed this issue by analyzing the cortical responses to spoken, written and signed sentences at the level of individual subjects. By applying a novel fingerprinting method based on the distributed pattern of brain activity, we identified a left-lateralized network composed by the superior temporal gyrus/sulcus (STG/STS), inferior frontal gyrus (IFG), precentral gyrus/sulcus (PCG/PCS), and supplementary motor area (SMA). In these regions, the local distributed activity pattern induced by any of the three language modalities can predict the activity pattern induced by the other two modalities, and such cross-modal prediction is individual-specific. The prediction is successful for speech-sign bilinguals across all possible modality pairs, but fails for monolinguals across sign-involved pairs. In comparison, conventional group-mean focused analysis detects shared cortical activations across modalities only in the STG, PCG/PCS and SMA, and the shared activations were found in two groups. This study reveals the core language system in the brain that is shared by spoken, written and signed language, and demonstrates that it is possible and desirable to utilize the information of individual differences for functional brain mapping.

## Introduction

Human languages are typically expressed by one of the three modalities: speech in the form of audio waves, text in the form of visual symbols, and sign in the form of visual-motor sequences. Despite remarkable differences in perceptual forms, the three types of language are structured and organized by similar linguistic rules in the domains of phonology, semantic and syntax (Sandler & Lillo-Martin, 2006). Where, along the neural path from sensory stimulation to ultimate comprehension, do the three processing streams converge for supramodal processing? Clarifying this issue can reveal the neural underpinnings of core language functions and provide important insights into the evolution of language (M. MacSweeney et al., 2008).

A number of studies have investigated the neural substrates that are shared by language processing in different modalities, yet the spatial extent of the observed supra-modal language network seems to vary from study to study (e.g.,Sakai et al., 2005; Lindenberg & Scheef, 2007; Weisberg et al., 2015). Among the many factors that potentially cause the inconsistency across studies, a widely-seen but uncontrollable factor is inter-subject variability. Subjects from different studies, and those participating in the same study, vary in language proficiency level, age of acquisition, conceptual space, and several other aspects. Due to those variabilities, the way how one perceives and interprets a given word or sentence, and the associated brain processes, differs markedly from other individuals (Buchweitz et al., 2009; Tanner & Van Hell, 2014; Nijhof & Willems, 2015).

The inter-individual variability has been conventionally treated as “noise” that hinders researchers to make inference about the general blueprint of functional brain, and been discarded via averaging the data across many subjects (Van Horn et al., 2008). Nevertheless, it is increasingly recognized that such variability should be taken as a source of information that is critical for a precise mapping between human cognition and the brain (Van Horn et al., 2008; Dubois & Adolphs, 2016; Seghier & Price, 2018). Indeed, recent empirical evidence has demonstrated that functional brain connectivity profiles (Finn et al., 2015), dynamic brain connectivity patterns (J. Liu et al., 2018), brain responses to tasks (Tavor et al., 2016) and whole-brain white matter architecture (Horien et al., 2019) are highly variable between subjects but highly consistent within individuals.

Grounded on an individual variability perspective, here we hypothesize, if language is processed at a certain cortical region independent of its form, an individual’s activity pattern at this region induced by one language modality should be able to predict his or her activity pattern induced by other modalities. Moreover, given that everyone is unique, the prediction across language modalities should be individual-specific. That is, the cross-modal prediction would be more effective within an identical individual compared to across different individuals.

To test the above hypothesis, we applied functional MRI to record the cortical responses of speech-sign bimodal bilinguals when they comprehended sentences in the forms of speech, text and sign respectively. Task-induced distributed activity patterns (DPAs) of brain were extracted and each subject’s DPAs induced by one modality were then utilized to distinguish the given subject from a group under another modality. To distinguish from the existing fingerprinting technique based on brain connectivities, we call this method “DPA-based fingerprinting”. If the DPA of a cortical region induced by one modality can be predicted by another modality and such prediction is individual-specific, the identification for individuals from a group will succeed across modalities. To establish the relevance of the supramodal network we may find to linguistic computations, we also applied this method to monolinguals who had no knowledge for sign language. It is expected that the DPA-based identification for the monolinguals will succeed across the spoken-written modalities, but will fail across signed-spoken and signed-written modalities due to their lack of linguistic knowledge for signs. To gain more insights into the nature of the supramodal network derived from this individual-focused approach, we also compared it with results derived from the group mean-focused conventional group activations analysis.

## Materials and Methods

### Participants

Fourteen bimodal bilinguals (3 males, aged 33 – 65 years) and 15 monolinguals without knowledge for sign language (3 males, aged 31– 67 years) were recruited. The bilinguals were native Chinese speakers, and acquired Chinese sign language (CSL) as their second language with high proficiency. As teachers in schools for the deaf, the bilinguals used CSL for at least 3.3 hours each day and had 12 – 40 years of CSL experience. The two groups were matched in age (*t*_(27)_ = 0.204, *p* = 0.84, two-tailed t-test) and education level (*t*_(27)_ = 0.144, *p* = 0.89, two-tailed t-test). All participants were right-handed according to the Edinburgh Handedness Inventory (Oldfield, 1971), and reported no neurologic or mental disorder. Written informed consent was obtained from all participants under a protocol approved by the Institutional Reviewer Board of Beijing Normal University. The same participants engaged in the sign language task were analyzed in our previous work addressing a different question (L. Liu et al., 2017).

### Experimental Design

During the fMRI scanning, 60 unrelated short declarative sentences were presented to every participant, with 20 in each modality. These sentences were adopted from previously published studies on sign and spoken language comprehension and were translated from English into Chinese (Neville et al., 1998; Mairéad MacSweeney et al., 2002; Mairéad MacSweeney et al., 2006). A part of the experimental materials and their English translations are provided in the Supplementary Material. A block design including four task blocks alternating with four baseline blocks was adopted. Each block lasted about 30 seconds during which five sentences were presented. For the spoken modality, audio recordings of sentences were played normally during task blocks, and played in reverse during baseline blocks. To avoid shared sensory inputs with other modalities, participants were required to have their eyes closed while listening to the speech. For the signed modality, silent videos of signed sentences were presented during task blocks, and videos showing the same signer standing still were presented during baseline blocks. For the written modality, written sentences and sequences of pseudowords were shown during the task and baseline blocks, respectively. Participants were told to comprehend those sentences, with no explicit response required. Examples of experimental stimuli are illustrated in Fig. 1A.

**Fig. 1.**
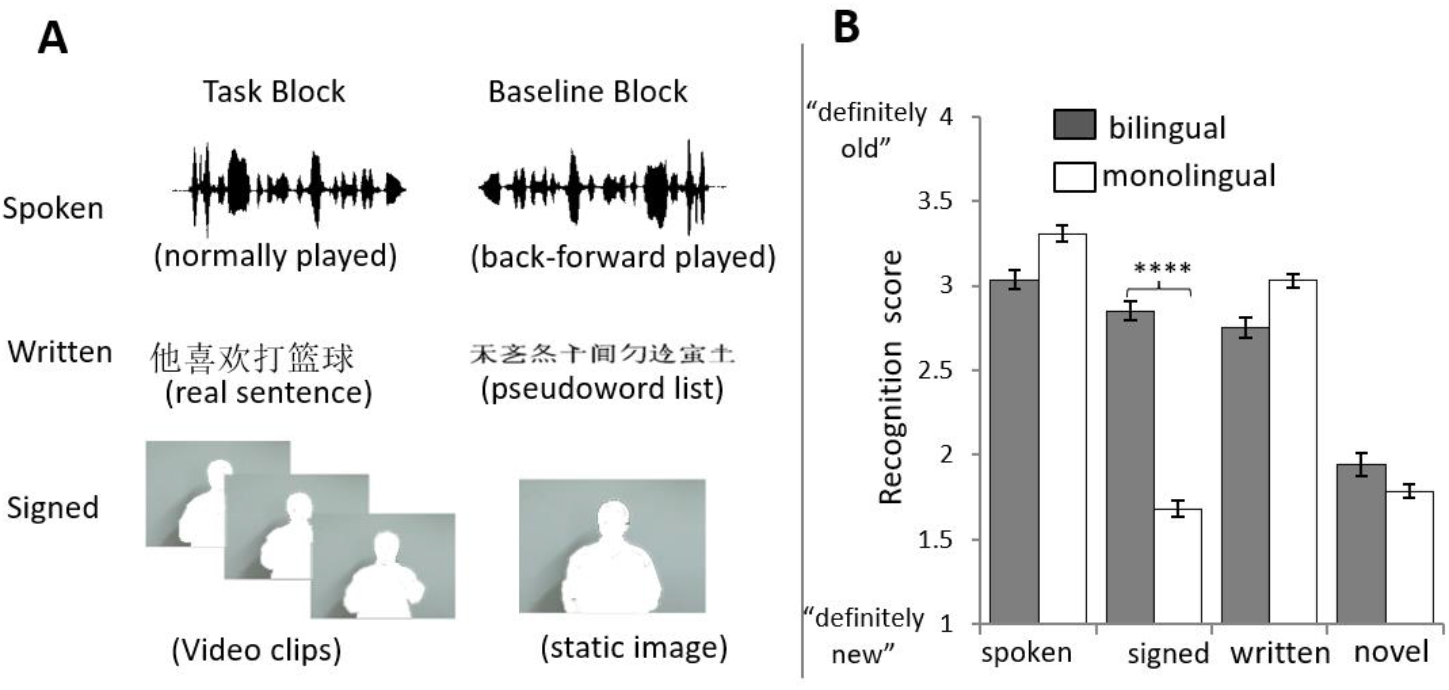
Examples of experimental stimuli (A) and behavioral performance (B). Bi-modal bilinguals and monolinguals were asked to comprehend spoken, written and signed sentences presented in different runs of fMRI scanning. In the post hoc behavioral test, the bilinguals performed better than monolinguals in the recognition of signed sentences. There was no significant group difference in the recognition of spoken or written sentences. **** *p* < 0.0005.

Following the scan, participants were given an unexpected recognition test in paper-pencil form. A total of 60 sentences were listed, half of which had been previously presented (10 for each modality) and half of which were novel. The order of sentences was randomized. Participants were asked to judge whether they had seen those sentences during the scanning session on a 4-point scale, with 1 defined as “definitely no (new)” and 4 defined as “definitely yes (old)”. We failed to record five subjects’ performance on the recognition test. Thus, the behavioral results were based on data from the remaining 24 participants.

### Image acquisition

Participants implemented the sentence comprehension task for the three modalities in three separate runs. The order of modalities was counterbalanced across participants. For each run, a total of 120 volumes were acquired in 6 min. Brain imaging data were collected with a 3T Siemens Trio Scanner at the MRI Center of Beijing Normal University. A gradient echo planar imaging (EPI) sequence was applied to functional image acquisition, with the following parameters: time repetition = 2000 ms, time echo = 30 ms, flip angle = 90°, filed of view (FOV) = 200 mm, matrix size = 64 × 64, slice number = 32 interleaved, and voxel size = 3.12 × 3.12 × 4.8 mm^3^. A MPRAGE sequence was adopted to acquire T1 structural images. Parameters were as following: time repetition = 2530 ms, time echo = 3.39 ms, flip = 7°, FOV = 256 mm, interleaved scan order, matrix size = 256 × 256, and voxel size = 1.0 × 1.0 ×1.33 mm^3^.

### Data preprocessing

Data preprocessing was performed using DPARSF (Yan & Zang, 2010) (http://rmfri.org/DPARSF), which integrates the preprocessing modules of Statistical Parametric Mapping (SPM12) (http://www.fil.ion.ucl.ac.uk/spm). The preprocessing included slice-timing correction, spatial realignment to correct for head motion, normalization to Montreal Neurological Institute (MNI) space based on individual’s T1 image segmentation, and resampling to 3 × 3 × 3 mm^3^ voxel size. No smoothing to the images was applied. One participant (bilingual) was excluded from further analysis due to excessive head motion (> 3 degree in rotation or 3 mm in translation) in the sign comprehension task.

### DPA-based fingerprinting analysis

#### Identification procedure

The analytical protocol is illustrated in Figure 2. First, task-induced brain activities were extracted for each subject using a general linear model (GLM) in SPM12. In this model, the sentence comprehension condition and the baseline condition plus six head-motion parameters were taken as regressors. Through this first-level analysis, we obtained a *t*-map for each subject and for each modality. Next, task-induced distributed pattern of activity (DPA) was obtained by using a searchlight method across the whole brain. At each voxel, a sphere containing 125 voxels centered on that voxel was generated, and *t*-values of the 125 voxels from the individual *t*-map were extracted and stored as the DPA of the central voxel. By this feature-extraction step, we created three datasets (corresponding to the three modalities) consisting of voxel-wise DPAs for all subjects in a group.

**Fig. 2.**
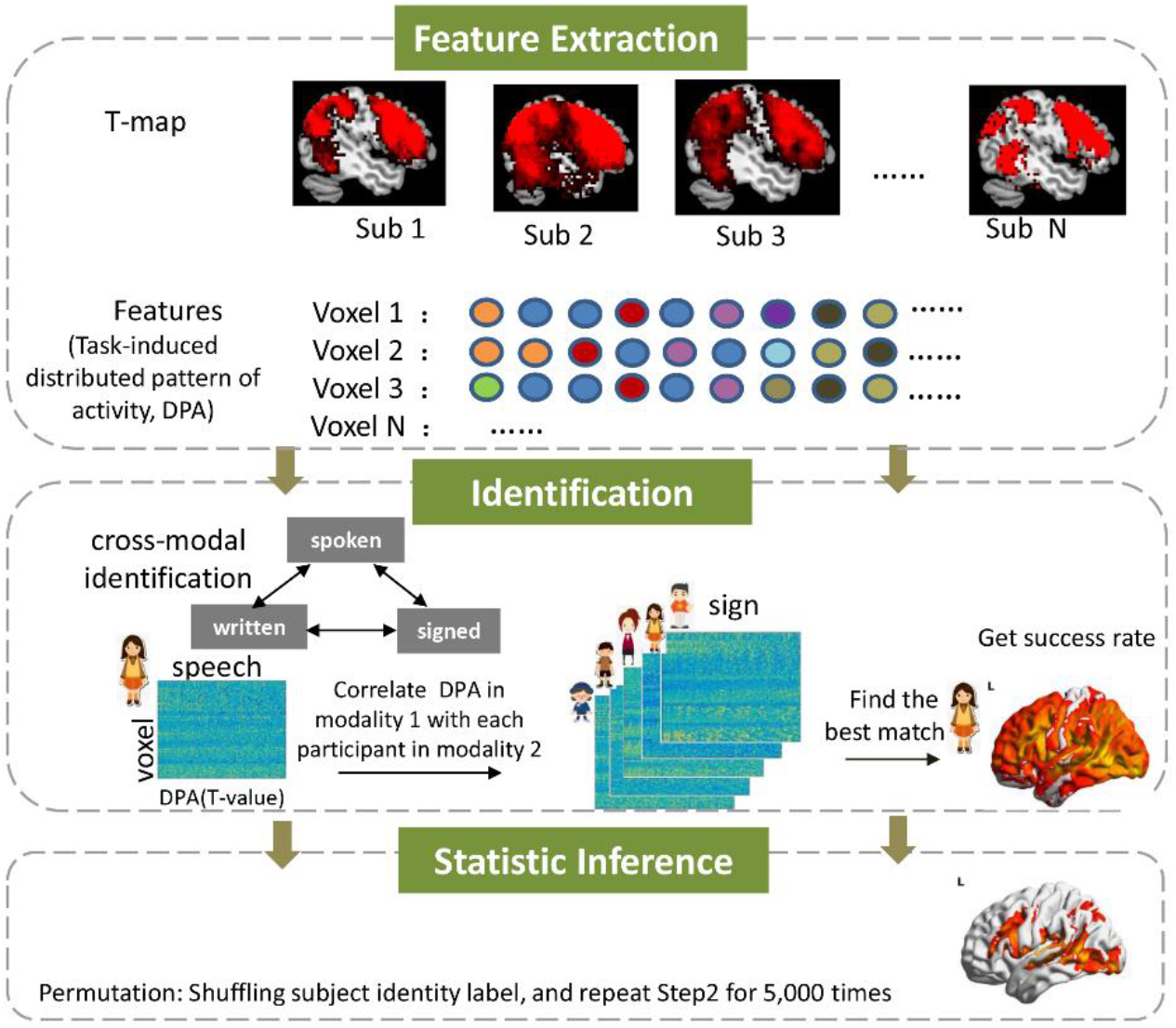
Schematic overview of the identification procedure. (1) At each voxel, a sphere containing 125 voxels centered on that voxel was generated, and *t*-values of the 125 voxels from the individual *t*-map (task > baseline) were extracted and stored as the DPA for the central voxel. (2) Identification for individuals across language modalities was made based on the similarity of a subject’s DPA induced by one modality to the DPA of every subject induced by another modality. If the one showing the highest correlation came from the same subject, the identification was successful. (3) To determine whether the success rate was statistically meaningful, a permutation test was conducted by randomly assigning participants’ identities of the target set and then performing the identification. These steps were iterated over all voxels in the brain, and an identification map showing voxel-wise success rates was generated for each pair of modalities.

The identification was performed for the bilingual and the monolingual groups separately. To test the robustness of findings, we also conducted the identification analysis by pooling bilinguals and monolinguals together into one group (see supplementary material). To predict a subject’s identity, similarity between the DPA of the given subject in one modality (base) and that of each subject in another modality (target) was assessed by using a Pearson’s correlation. If the two DPAs showing the strongest correlation came from the same subject, the identification was successful, otherwise it failed. This procedure was repeated across all subjects and all possible combinations of modalities (spoken-to-signed, signed-to-spoken, spoken-to-written, written-to-spoken, signed-to-written, written-to-signed). The identification was iterated across all voxels in the brain.

### Success rate and statistical analysis

While activity patterns vary, we assume the anatomic locations of the core language system in brain are grossly fixed across subjects (despite it could also vary to some degree). On the basis of this premise, we calculated for each voxel the success rate of identification as the percentage of subjects whose identities were correctly predicted out of the total number of subjects in the group (Finn et al., 2015), producing a whole-brain map of identification rate for each pair of modalities. The success rate of a voxel reflected comprehensively the degree of cross-modal stability and individualization in its activity pattern, as well as the degree of consistency in exhibiting such features across subjects.

To assess whether the obtained success rate was significantly above chance, nonparametric permutation tests were performed. Specifically, one modality was randomly chosen to serve as the base set, and another was randomly chosen from the rest of modalities to serve as the target set. Then, subjects’ identities of the target set were randomly assigned and the identification was performed. This procedure was repeated 5,000 times for every voxel across the whole brain to create null distributions. Empirical p-values were obtained by calculating the fraction of samples in the null distribution whose identification accuracies were higher than the observed one. Statistical significance was set to FDR corrected *p* < 0.005 and a cluster size > 30 voxels.

To detect those cortical regions with successful identification across all modalities, a conjunction map was generated through an intersection across six thresholded identification maps (corresponding to the six pairs of modalities), and the mean of success rate across these thresholded maps was computed. Conjunction maps across two modalities (e.g., spoken-to-signed and signed-to-spoken) were also generated.

### Identification using the activity pattern of a network and the whole brain

In the above DPA-based fingerprinting analysis, we had used only the information from a small volume of the brain (125 voxels with a volume smaller than 4 cm^3^) as features to characterize an individual brain in order to achieve a high spatial resolution. However, prior fingerprinting studies typically use the connectivity profiles of a large network or of the whole brain as features (e.g., 268 nodes, 35,778 edges) (Finn et al., 2015; Vanderwal et al., 2017). To make the findings from this study comparable with findings from prior work, we also performed the identification using features extracted from a network and from the whole brain. Here, we examined specifically the supramodal network derived from the DPA-based fingerprinting analysis. The *t*-values of all voxels within the network (or the whole brain) were extracted from an individual *t*-map to create an activity pattern for each subject and each modality. During the creation of whole-brain activity pattern, a mask generated by a SPM second-level analysis for group activation (reported below) was applied to exclude voxels outside the brain. The identification procedure applied was the same as described above.

### Conventional activation analysis

To gain insights into the nature of the supramodal network we may find via the novel DPA-based fingerprinting analysis, we compared it with the results from conventional activation analysis. For the conventional activation analysis, the preprocessed functional images were further spatially smoothed using a Gaussian kernel with 7 mm full-width at half maximum. Applying the same GLM as described above, we first examined the effect of task versus baseline for individual subjects. Next, a second-level analysis (one-sample t-test) was conducted to examine the group-mean effect of cortical activation. In order to identify those anatomic loci being significantly activated by language comprehension regardless of the modality, the group *t*-maps for the three modalities were thresholded first, and then a conjunction map was generated by computing the intersection across the three thresholded *t*-maps.

### Contribution of within-subject stability and between-subject variability to the identification performance

The ability of a cortical region to accurately identify individuals from a group was mainly determined by 1): how stable was its activity pattern within an individual across modalities (within-subject stability); and 2): how variable was its activity pattern across different subjects (between-subject variability). To understand why some cortical regions identified individuals successfully while other regions failed, we measured the within-subject stability and between-subject variability across all voxels in the brain, and then assessed the respective contributions of the two factors to the identification performance. Here, voxels with greater accuracy than the permutated ones were labeled as “success” and otherwise as “failure”. A multiple logistic regression model was applied to predict the identification performance. This analysis was conducted using the data and the identification results for bilinguals across as a single pair of modalities (reported is the identification from speech to sign). Before entering the logistic model, the measurements for within-subject stability and between-subject variability were normalized across voxels using *z*-score transformation. The method for quantifying within-subject stability and between-subject variability is provided in the supplementary material.

### Potential contribution of intrinsic process to the identification

Recent work has demonstrated that individuals can be distinguished from a group based on their task-free, intrinsic functional connectivity profiles (Finn et al., 2015; Tavor et al., 2016; Horien et al., 2019). Thus, the individual-specific activity pattern we aimed to identify could arise from individual’s unique intrinsic cognitive process, rather than from the cognitive process related to the ongoing task. Since the task-free intrinsic process and the task-related cognitive process are typically associated respectively with a decrease and an increase in BOLD responses relative to a low-level baseline (Raichle et al., 2001; Harrison et al., 2007), we re-analyzed the data using only task-positive activities and only task-negative activities, respectively. For the identification using only task-positive activities, voxels with *t*-values < 0 were set to zero when creating the searchlight pattern. For that using only task-negative activities, voxels with *t*-values > 0 were set to zero.

## Results

### Behavioral results

On the old/new recognition test, the spoken and written sentences were scored higher (indicate more likely to be “old”) than novel sentences by both groups. Significant group difference was only found in the recognition for signed sentences (bilingual > monolingual, *t*_(23)_ = 5.15, *p* < 10^−4^, two-tailed t-test), but not for spoken, written or novel sentences (all *p* values > 0.20). The bilinguals scored signed sentences higher than novel sentences (*t*_(10)_ = 5.56, *p* < 10^−3^, paired t-test), while monolinguals scored them equally (*t*_(12)_ = −0.99, *p* = 0.34, paired *t*-test) (Fig. 1B). These results suggest that all subjects comprehended those written and spoken sentences, but only the bilinguals comprehended the signed sentences.

### A left-lateralized network allowing successful identification of bilinguals across modalities

We hypothesize that, if language is processed by a certain cortical region independent of its form, the activity pattern of this region induced by one modality will be able to predict the activity pattern induced by any of the other two modalities, and the cross-modal prediction will be individual-specific. In line with this hypothesis, the DPA-based fingerprinting analysis detected a set of cortical regions wherein the activity pattern induced by one modality was similar to that induced by another modality, and the degree of cross-modal similarity was greater within an identical subject than between different subjects. The stable and individualized activity pattern across modalities allowed us to identify an individual among a group of subjects. Anatomic loci with success rates of identification greater than chance across all six pairs of modalities were dominantly located in the left hemisphere, encompassing the left posterior STG/STS extending to MTG, precentral gyrus/sulcus (PCG/PCS), IFG, and bilateral supplementary motor area (SMA) (Fig. 3A) (threshold: p < 0.005, FDR corrected for whole-brain multiple comparisons). Cortical maps displaying cross-modal similarity within and between individuals are provided in the supplementary material (Fig. S1). A similar supramodal network was derived from the fingerprinting analysis wherein the two groups were pooled together during the identification procedure (Fig. S2).

**Fig. 3.**
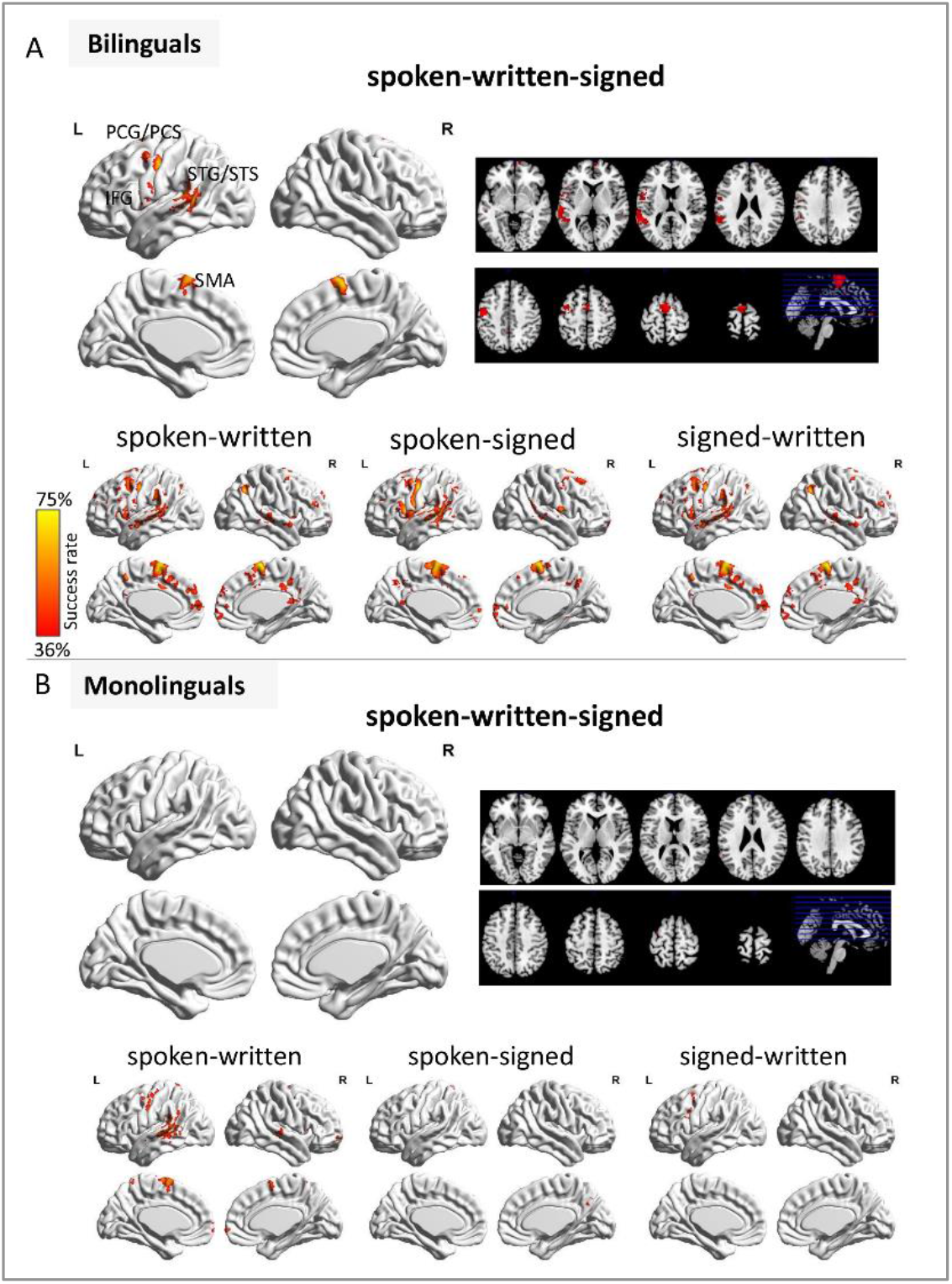
The supramodal network allowing successful identification for individuals across language modalities. We attempted to identify an individual among a group of subjects engaged in one modality based on his or her activity pattern induced by another modality. This method detected a set of cortical regions that allowed the identification for bilinguals across all six pairs of modalities (denoted by “spoken-written-signed”), with success rates greater than that obtained from the permutation tests (A). In comparison, no voxel across the whole brain allowed successful identification for monolinguals across all six pairs. This was due to the identification failure specific to the signed-involved pairs (B). Thresholded: p < 0.005, FDR corrected for whole-brain multiple comparisons. The 3D surface visualizations of the results are implemented using the BrainNet Viewer (www.nitrc.org/projects/bnv).

### Identification for monolinguals fails across sign-involved modality pairs

To determine whether the individual distinguishing property of the supramodal network revealed above was driven by language-specific processing or instead by other non-linguistic processes (such as working memory, attention, or task-unrelated intrinsic process), we applied the fingerprinting analysis to monolinguals who did not understand sign language. In contrast to the bilinguals, there was no voxel across the whole brain that allowed successful identification for the monolinguals across all six pairs of modalities. Further inspection revealed that, the left STG, SMA and PCG/PCS allowed the identification for monolinguals across the spoken-written pair. However, the identification across signed-spoken and signed-written pairs largely failed (Fig. 3B). The specificity of identification failure for monolinguals to the sign-involved pairs suggests that the individual distinguishing property of the supramodal network was driven by language-specific processing.

### Comparison with the results from conventional activation analysis

To gain more insights into the nature of the supramodal network with individual distinguishing property, we compared it with the results from conventional activation analysis. When the same threshold as used in the fingerprinting analysis (FDR corrected p < 0.005) was applied to detect significant cortical activations, there was no overlap in cortical activation across the three modalities. When applying a more lenient threshold (uncorrected p < 0.01, cluster size ≥ 30 voxels), we found overlap in cortical activation across the three modalities in the left PCG, left STS/MTG and bilateral SMA in the bilingual group (Fig. 4A). Nevertheless, with this lenient threshold, activation overlap across the three modalities was also observed in the monolingual group in the left PCG and left STS/MTG.

**Fig. 4.**
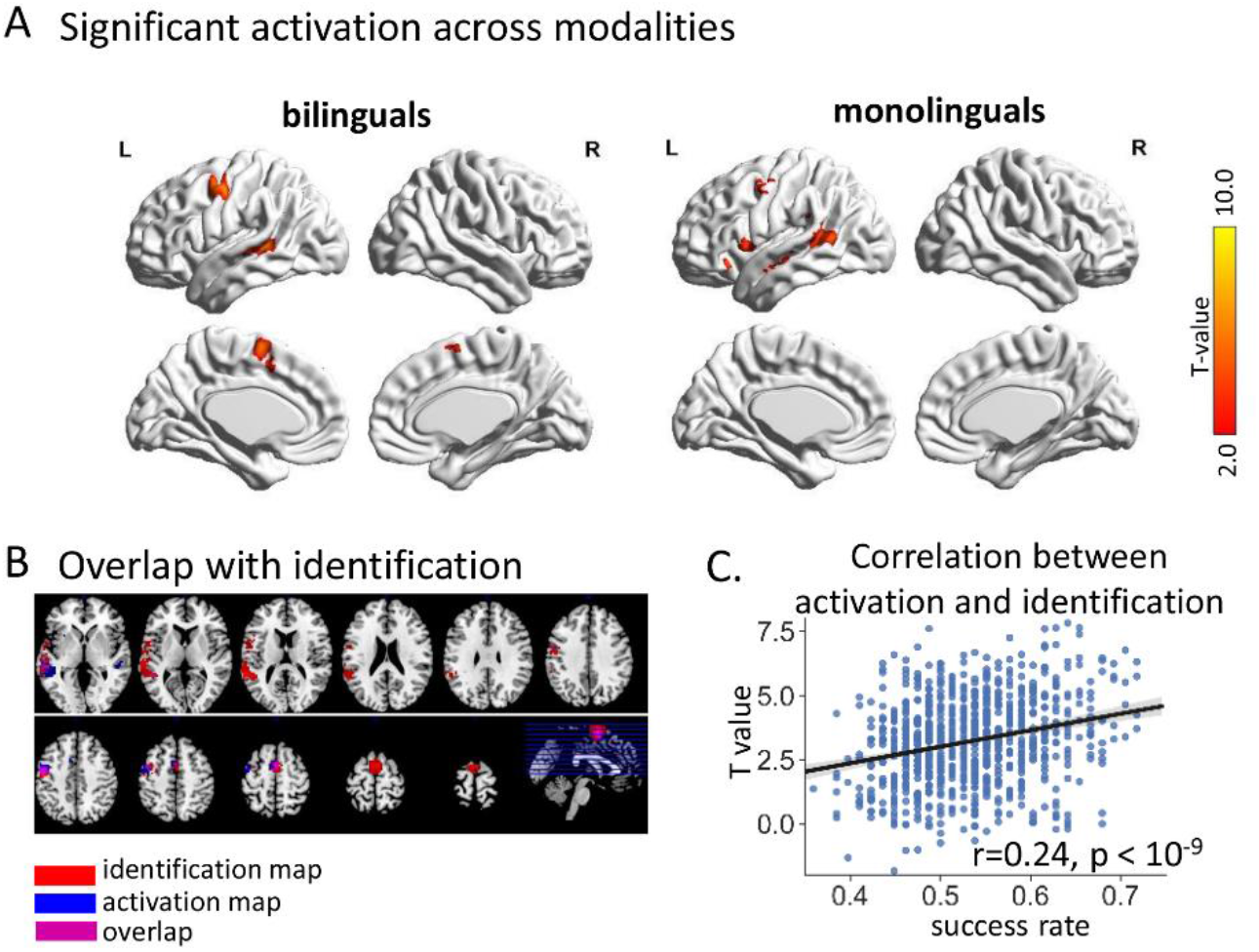
Comparison with the results from conventional activation analysis. (A) Cortical regions consistently activated across the three modalities. Note, modality-consistent cortical activations were found in both groups. Threshold: uncorrected *p* < 0.01 combined with a cluster size ≥ 30 voxels. (B) Spatial overlap between the supramodal network derived from the fingerprinting analysis and the modality-consistent activation map (for bilinguals). (C) Significant positive correlation between success rate and activation magnitude across voxels (N = 887) within the supramodal network.

Compared to the activation map, the supramodal network revealed by the fingerprinting analysis was spatially more extensive (337 versus 886 voxels). Among those voxels with significant activation for all three modalities, 53.5% of them fell into the supramodal network derived from the fingerprinting analysis (Fig. 4B). Interestingly, the success rate (modality-averaged) of identification was positively correlated with the activation amplitude (modality-averaged group *t*-values) across voxels within the supramodal network (Pearson’s r = 0.26, p < 10^−9^, N = 886 voxels) (Fig. 4C). This means that the stronger a voxel was activated during language comprehension, the more accurate it was in identifying individuals from a group. These findings indicate that the supramodal network emerged from the fingerprinting analysis due to its engagement in the ongoing task.

### The activity pattern of the supramodal network acts as a “fingerprint”

For the supramodal network derived from the DPA-based fingerprinting analysis, despite the success rate of identification (mean: 53.1%; range: 36%–72% across voxels within this network) was greater than chance, it was not as high as that reported in previous connectome-based fingerprinting studies (about 54% to 99% in Finn et al. (2015)). This might be because the size of feature (a searchlight consisting of 125 voxels) we had used for the identification was too small as compared to that in previous studies (e.g., 35,778 features in Finn et al. (2015)). To test this possibility, we performed the identification using the activity pattern consisting of all voxels within the supramodal network (N = 886 voxels). Indeed, the average identification accuracy for bilinguals rose to 91.3 % (range: 76.9% – 100% across modality pairs), suggesting the activity pattern of the supramodal network can act as a “fingerprint” feature to identify individuals. For the monolinguals, the identification rate increased to 76.7% for the spoken-written pair, but was still low for the sign-involved pairs (mean: 54.9%, range: 53.3%–60%). Chi-square tests (one-tailed) showed that the identification for bilinguals was significantly more successful than that for monolinguals, especially on the sign-involved pairs (Fig. 5A). To give an impression about this “fingerprint”, we display the network activity pattern from three representative subjects (Fig. 5B). Consistent findings were obtained when the two groups were pooled together during the identification procedure (Fig. S3).

**Fig. 5.**
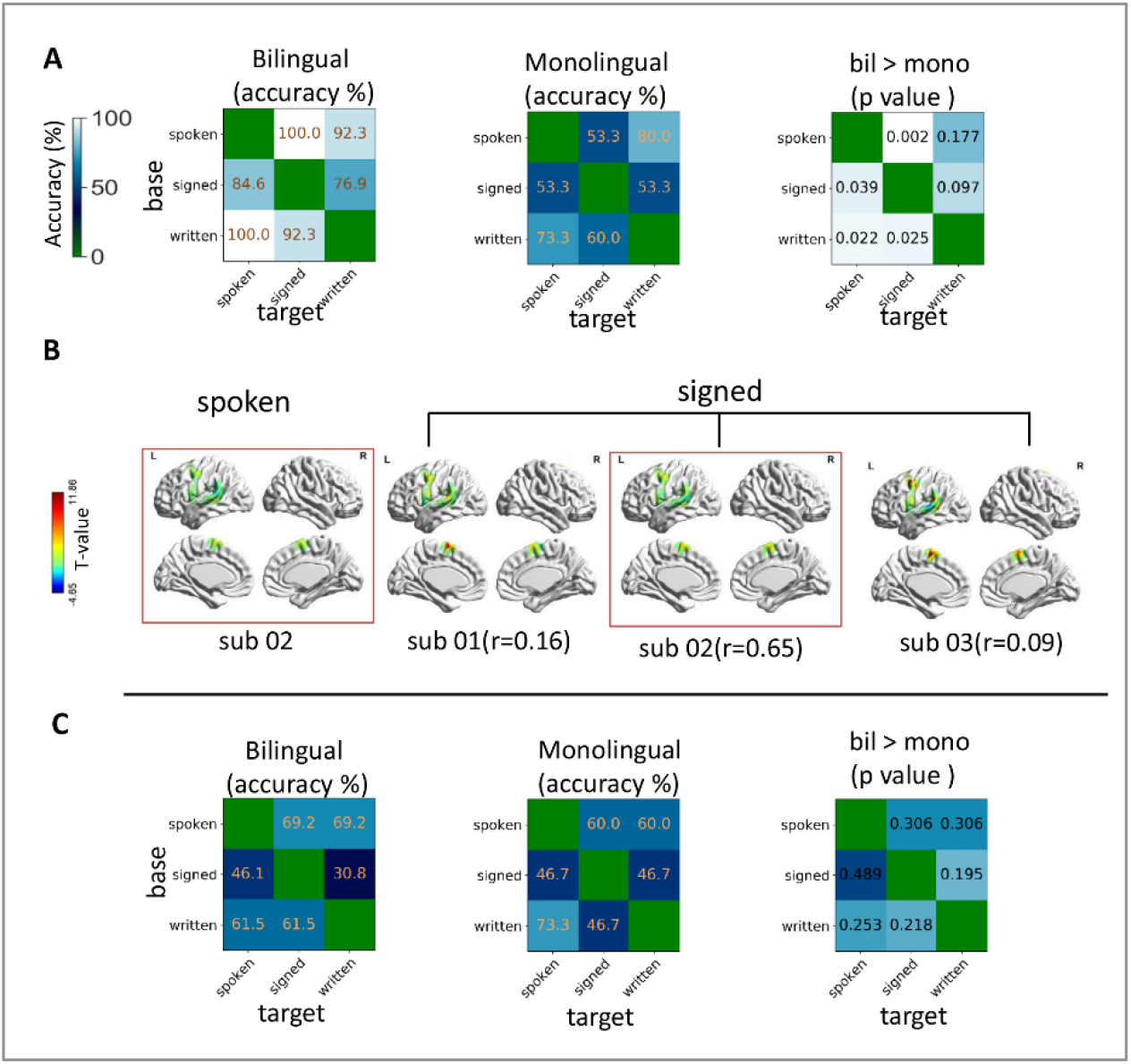
The activity pattern of the supramodal network acts as a “fingerprint”. (A) Identification using the activity pattern consisting of all voxels in the supramodal network (N = 886) yielded high success rates. The success rate for bilinguals was significantly higher than that for monolinguals, especially on sign-involved pairs. (B) The activity pattern of a representative subject in the spoken modality (base data) and those of her own and another two subjects in the signed modality (target data). The value in the bracket indicates the degree of similarity in activity patterns between the training and target data. (C) Identification accuracy was not improved by using the activity pattern consisting of all voxels in the brain (N = 59, 608), suggesting feature quality (the relevance to the task) is essential for successful identification.

To determine whether the increase in identification accuracy is simply due to the increase in feature size or depends also on the quality of the selected feature, we performed the identification analysis using an activity pattern consisting of all voxels in the whole brain (N = 59, 608 voxels). The identification (mean success rate for bilinguals: 56.4%; range: 30.9% – 69.2%) was not better than that obtained from the DPA-based identification. Moreover, the significant group differences in identification rates disappeared (Fig. 5C). These results suggest that the success in identifying individuals across modalities relies on both the quality (here refers to relevance to the task) and the size of feature.

### The contribution of within-subject stability and between-subject variability

To understand why some cortical regions identified individuals successfully across modalities while other regions failed, we built a multiple logistic regression model which took within-subject stability and between-subject variability as predictors and voxel-wise identification performance (categorized as “success” or “failure”) as an outcome. The results showed that both variables made significant positive contributions to the outcome (for within-subject stability: standardized *β* = 2.59, *p* < 10^−10^; for between-subject variability: standardized *β* = 0.83, *p* < 10^−10^; tested with the identification from speech to sign). This means that voxels with higher degrees of within-subject stability and between-subject variability were more likely to succeed in the identification. This trend is replicable by other modality pairs. Scatter plots showing the within-subject stability against between-subject variability for three modality pairs are provided in supplementary figure S5.

### The supramodal network was reproducible using task-positive activities but disappeared using task-negative activities

To assess the potential contribution of intrinsic process to individual identification, we conducted the fingerprinting analysis using only task-negative activities. This analysis yielded only a few voxels located in the medial frontal cortex that allowed successful identification for individuals across modalities. By comparison, the analysis using only task-positive activities yielded a similar identification map (Fig. S6) as that resulted from the analysis using both task-positive and task-negative activities. These findings suggest that the revealed supramodal network arose from the neural processes responsible to the ongoing language task, rather than from task-unrelated intrinsic processes.

## Discussion

Where is human language processed in the brain independent of its form? The understanding of this issue has mainly relied on neuroscientific evidence that examined only those brain processes that are common and universal across individuals. Grounded on an individual variability perspective, we applied a novel DPA-based fingerprinting approach to identify the neural correlates supporting supramodal linguistic processing. The analyses revealed a left-lateralized network wherein the DPAs induced by spoken, written and signed languages were highly variable across subjects but highly stable within subjects, allowing successful identification for individuals among a pool of subjects. The identification for bimodal bilinguals was successful across all possible modality pairs, but failed for monolinguals (without knowledge for sign language) across sign-involved pairs.

### The supramodal network is locked to linguistic computations

Three sets of results suggest that the supramodal network identified by the DPA-based fingerprinting analysis arose from linguistic computations shared by spoken, written and signed language, rather than from task-unrelated intrinsic or non-neuronal processes. First, the supramodal network allowed the identification for bilinguals across all six pairs of modalities, but failed in the identification for monolinguals who did not understand sign language. Importantly, the identification failure for monolinguals was specific to sign-involved pairs (signed-spoken and signed-written pairs), suggesting linguistic computations played a critical role in determining the individual distinguishing property of the supramodal network. Second, this network was reproducible using task-positive activities but disappeared using task-negative activities. This finding largely rules out the potential contribution from the intrinsic or non-neuronal processes. Third, the success rate of individual identification was positively correlated with the amplitude of group-level activation across voxels within the supramodal network, suggesting that the individual’ distinguishing feature of the supramodal network is dependent on its engagement in the ongoing task.

### Potential application of the DPA-based fingerprinting technique

Compared with the results obtained from the conventional activation analysis, the supramodal language network derived from the DPA-based fingerprinting analysis was spatially more extensive and statistically more robust. Significant cortical activations were detected in the left PCG, left STS/MTG and bilateral SMA for all three modalities. These cortical regions were also detected by the fingerprinting analysis. The fingerprinting analysis further detected the left pSTG/STS and IFG. Note, the threshold applied to the activation map (uncorrected p < 0.01) was more lenient than that applied to the identification map (FDR corrected p < 0.005). These findings indicate that the DPA-based fingerprinting analysis might be more sensitive than the conventional activation analysis to detect those neural substrates important for the ongoing task.

Current fingerprinting techniques typically use whole-brain or network connectivity profiles to distinguish individuals across different tasks and states, which capture those reliable individual differences associated with task-independent intrinsic process (Finn et al., 2015; Vanderwal et al., 2017; Horien et al., 2019). Differing from this technique, the DPA-based fingerprinting captures reliable individual differences driven by task-induced cognitive processing, and can localize these effects to the level of a focal voxel. The ability to identify reliable individual differences in task-induced brain activities with a high spatial resolution is important, since tasks can bring out meaningful variability across subjects and amplify individual differences in brain signals beyond what can be measured at rest. The DPA-based fingerprinting method thus provides a useful tool for future studies aiming to link individual differences in cognition and behaviors to individual differences in brain activities induced by relevant tasks.

### Limitations and future direction

The current study has two limitations. First, compared to those connectome-based fingerprinting studies, the current study had a smaller sample size, which might inflate the identification accuracy. However, we note that the identification procedure applied to the two groups separately (13 and 15 subjects) and that applied to the two groups polled together (28 subjects, results provided in the supplementary material) yielded quite similar results: the anatomic location of brain voxels with distinguishing property was consistent between the two approaches, and the identification accuracy for bilingual individuals based on the network activity pattern changed little (76.9%-100% for the smaller sample size; 76.9%-92.3% for the larger sample size). It is worthwhile to examine whether the individual-focused fingerprinting technique is more robust to sample size variations than group mean-focused analysis. Second, despite the group comparison and two additional analyses have established a tight link between the supramodal network’s distinguishing property and linguistic computations, it is unclear which components of linguistic computations makes the activity pattern of this network individually distinct.

## Conclusion

Applying a novel DPA-based fingerprinting method, the current study reveals a supramodal network, composed by IFG, pSTG/STS, PrCG/PrCS and SMA with left-hemisphere dominance, which is reliably and uniquely recruited by individuals to comprehend spoken, written and signed sentences. These findings provide new insights into the neural underpinnings of core language functions. Furthermore, this study demonstrates that it is possible and desirable to utilize the information of individual difference for functional brain mapping. Future studies can apply the DPA-based fingerprinting technique to examine how an individual brain is activated by a targeted task in a unique way, and use the individualized brain activation to explain or predict behavioral phenotypes.

## Acknowledgments

This work was supported by grants from the National Natural Science Foundation of China (NSFC: 31971036, 31900802), China Postdoctoral Science Foundation (2019M653248) and the Open Research Fund of the State Key Laboratory of Cognitive Neuroscience and Learning of Beijing Normal University (CNLYB1803).

