## supplementary material for "Identifying a supramodal language network in human brain with individual fingerprint"

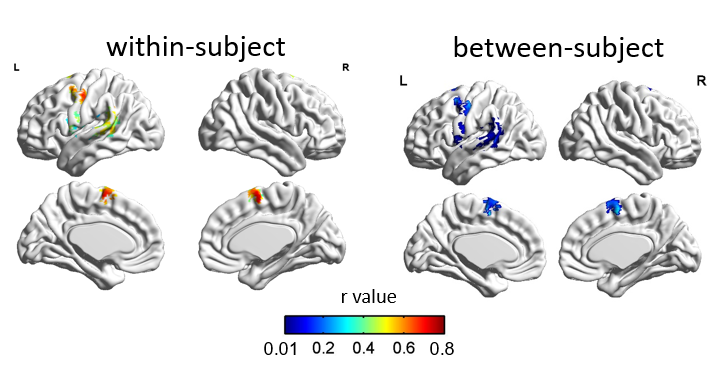

**Fig. S1.** Within- and between-subject similarity in distributed activity patterns across modalities. Displayed are the similarity measurements averaged across subjects (from the bilingual group) and across all six pairs of modalities. A detailed description for the assessment of within- and between-subject similarity is provided in the session “The quantification for within-subject stability and between-subject variability” below.

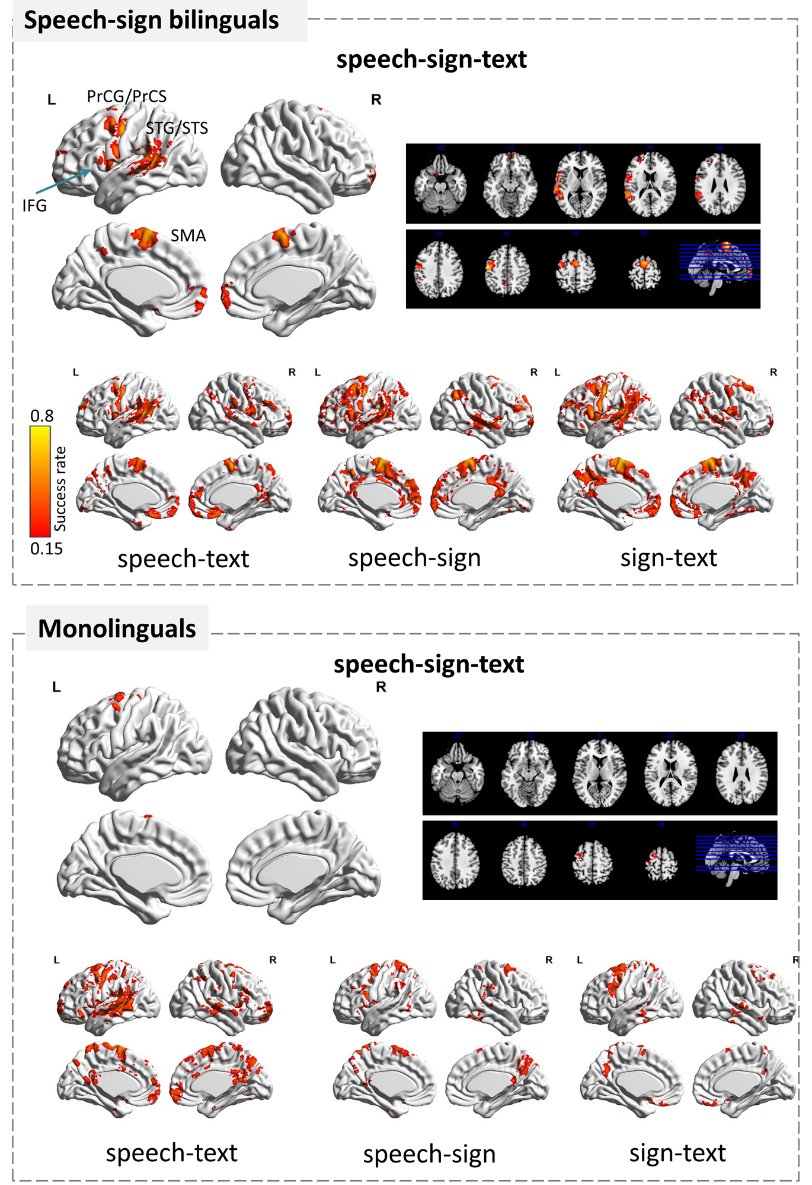

**Fig. S2.** Identification map obtained from the DPA-based fingerprinting analysis where bilinguals and monolinguals were pooled into one group. Here, a success rate was calculated as the percentage of subjects whose identities were correctly predicted out of the total number of subjects in each subgroup. Only voxels whose identification accuracies were higher than the best performance from 10,000 permutations were considered significant. The results were consistent with those obtained from the identification analysis conducted for the bilingual group and monolingual group separately.

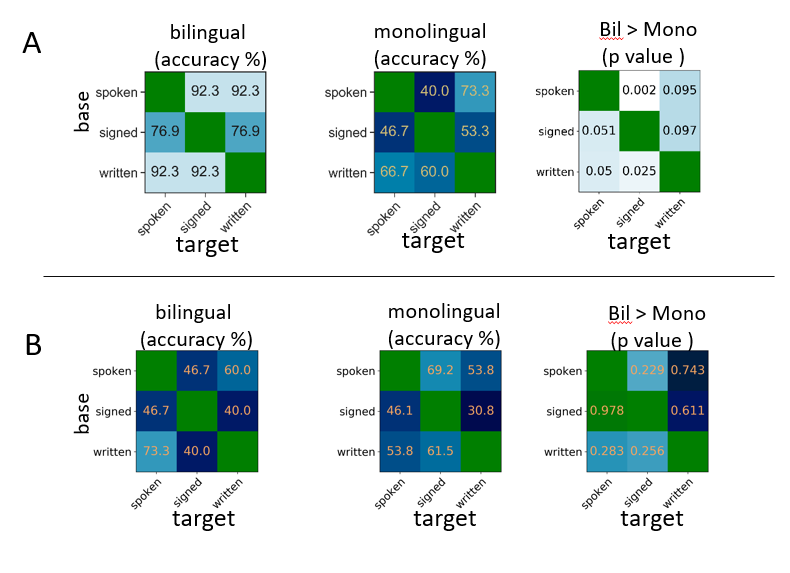

**Fig. S3.** Identification using the activity pattern of the supramodal network (A) and the whole brain (B). During the identification procedure, bilinguals and monolinguals were pooled into one group.

**The quantification for within-subject stability and between-subject variability**

To quantify within-subject stability (similarity), we first computed a Pearson’s correlation of the DPAs between two modalities for each subject. The r values were converted to z values by Fisher’s *r-z* transformation to improve the normality of the distribution of correlation values, and were then averaged across subjects. The averaged z values were then inverse transformed (z-to-r) to produce average r-values in the plots. The between-subject variability is defined as the between-subject similarity subtracted by one. To assess between-subject similarity, we first computed the mean correlation coefficient between a subject’s DPAs in one modality and the DPA of the rest of subjects in another modality, and then averaged the correlation value across subjects. Figure S4 illustrates the assessment of within- and between-subject similarity for one subject.

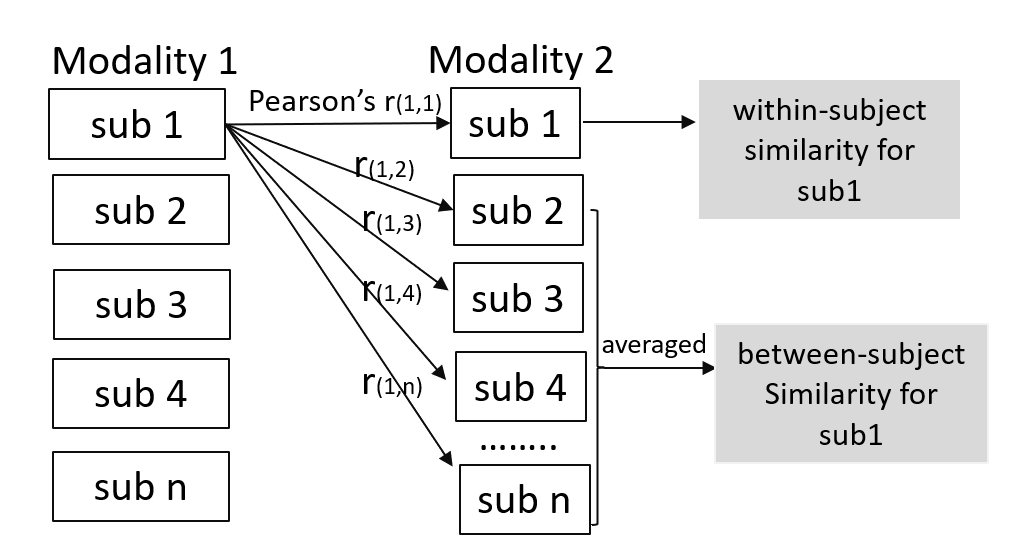

**Fig. S4.** The assessment of within- and between-subject similarity across modalities for one subject.

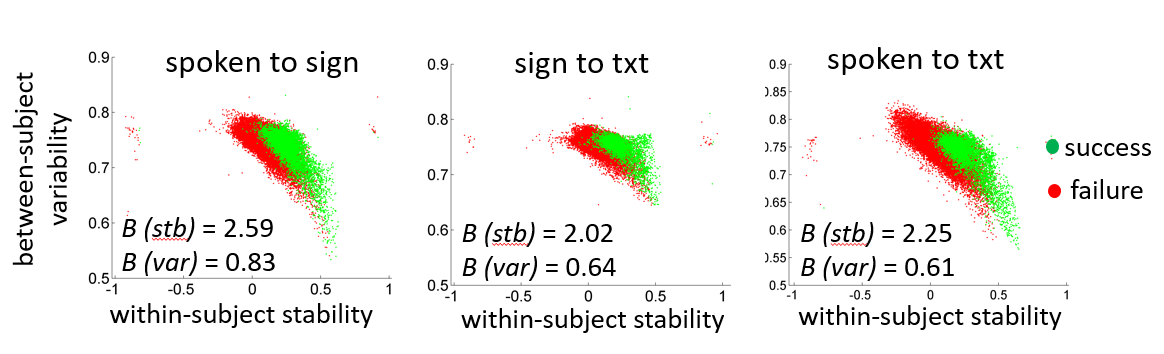

**Fig. S5.** Scatter plot showing within-subject stability against between-subject variability in DPAs for all voxels in the brain (N = 58,885 voxels). Both factors had a positive role in determining a voxel’s success in distinguishing individuals from a group of subjects. Note, among voxels with comparable between-subject variability, those showing a higher degree of within-subject stability were more likely to succeed in the identification. Among voxels with comparable within-subject stability, those showing a higher degree of between-subject variability were more likely to succeed.

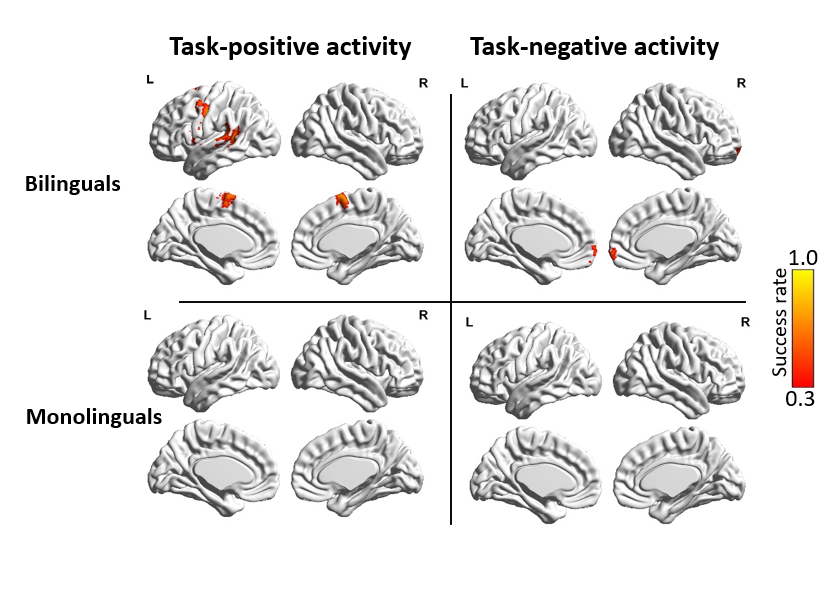

**Fig. S6.** Identification for individuals using only task-positive and only task-negative activities. The supramodal network yielded by the analysis using only task-positive activities is consistent with that yielded by the analysis using both positive and negative activities (shown in the main text). However, this network disappeared in the identification using only task-negative activities.

Table 1. A part of the sentences presented to subjects during fMRI scanning.

| Chinese | English translations |
| --- | --- |
| 组一:  1. 我很喜欢爸爸的那辆汽车 | Block 1:  1. I like Dad's car very much |
| 2. 我把帽子从箱子里取出来 | 2. I took the hat out of the box. |
| 3. 明天的语文考试我准备得不好 | 3. I didn't prepare well for the language test tomorrow. |
| 4. 他们没有决定去哪玩 | 4. They haven’t decided where to have fun. |
| 5. 我收到了妈妈寄来的一封信 | 5. I received a letter from my mother. |
| 组二: | Block 2: |
| 1. 来自聋校的三个女孩在打篮球 | 1. Three girls from the deaf school are playing basketball. |
| 2. 妈妈递给我一杯水 | 2. Mom handed me a glass of water. |
| 3. 钥匙放在左边第二个抽屉里 | 3. The key is in the second drawer on the left |
| 4. 我很喜欢阿姨的那条项链 | 4. I like Auntie's necklace very much |
| 5. 你可以要一个苹果或一个香蕉 | 5. You can ask for an apple for a banana. |
